# Establishing Novel Doxorubicin-Loaded Polysaccharide Hydrogel for Controlled Drug Delivery for Treatment of Pediatric Brain Tumors

**DOI:** 10.1101/2022.05.23.493140

**Authors:** Jenny P. Patel, Alissa Hendricks-Wenger, Carli Stewart, Kassidy Boone, Naydia Futtrell-Peoples, Lyndon Kennedy, Elizabeth D. Barker

**Affiliations:** Department of Mechanical, Aerospace, and Biomedical Engineering, The University of Tennessee at Knoxville, Knoxville, Tennessee; Lincoln Memorial University - DeBusk College of Osteopathic Medicine, Knoxville, Tennessee

**Author notes:** **Corresponding author:** Elizabeth D. Barker.

**Keywords:** Doxorubicin, hydrogel, amylopectin, medulloblastoma, local drug delivery, pediatric

## Abstract

According to the National Cancer Institute, of the more than 10 million cancer survivors alive in the United States at least 270,000 were originally diagnosed under the age of 21. While the 5-year survival rates for most childhood cancers appear very promising, the long-term survival rates are still very dismal. There is significant long-term morbidity and mortality associated with treatment of childhood cancer, and the risk of these effects continues to increase years after completion of therapy. Among childhood cancer survivors the cumulative incidence of a chronic health condition is 73.4% 30 years after the original cancer diagnosis, with a cumulative incidence of 42.4% for severe, disabling, life-threatening, or death due to a chronic condition caused by the chemotherapy used to treat the initial malignancy. Brain tumors are the most prevalent solid tumor diagnosed in children, and account for 20 percent of all childhood cancer deaths. The efficacy of all chemotherapy agents can be limited by their toxicity, their instability, and their ability to be formulated into practical drug products for use in the clinical setting To address this gap, our group has developed a novel carbohydrate-based hydrogel, Amygel, that is capable of being loaded with drugs and injected directly into the site of disease. Local drug delivery using Amygel has potential to improve childhood cancer treatment outcomes and prevent the devastating effects of systemic chemotherapy exposure. Development of Amygel for clinical use has three focus areas including: increasing drug concentration at the target site; improving chemotherapy penetration through tumor tissue, and; establishing chemotherapy dosage forms for pediatric use. For this study, we formulated Amygel with dimethyl sulfoxide and integrated the chemotherapy doxorubicin (DOX). High-performance liquid chromatography (HPLC) was used to confirm the quality of DOX after hydrogel synthesis, rheology and syringability tests to characterize the mechanical properties, and performed an *in vitro* cytotoxicity test against the pediatric medulloblastoma cell line DAOY. On HPLC, we found that after integrating DOX into the Amygel matrix the drug maintained a strong band on the chromatograph at the same point with the same intensity as the control free drug, indicating there were no changes in the structural properties of DOX. The mechanical tests showed that there was a proportionate increase in the storage modulus of the drug-loaded hydrogels as the concentration of amylopectin increased from 3 wt% to 20 wt%, but even at 20 wt% the hydrogel remained below the medical standard for injectables that the burst force should not exceed 40 N and the sliding force below 20 N. Correlating with the rheology findings, as the concentration of amylopectin increased, and therefore the strength of the hydrogel, there was an increase in the magnitude of force required for gel injection. These mechanical studies additionally provide evidence that the mechanical stability of the gel is not dampened by the incorporation of DOX. Drug release and cytotoxicity studies demonstrated a sufficient release of DOX from the hydrogels, and that the DOX released was able to achieve significant (p<0.01) cell death.

## Introduction

Globally, over 300,000 new cancers are diagnosed annually in patients younger than 19 years old, with the most common being central nervous system tumors [1]. In the past fifty years, overall survival rates have significantly improved, but long-term survivorship, quality of life, and improving treatment outcomes for aggressive tumor subtypes have remained challenging issues in pediatric oncology. Brain tumors are the most common solid tumor diagnosed in pediatric patients and medulloblastoma is the most common pediatric brain tumor. Chemotherapies administered orally and intravenously are part of the standard of care treatment for most pediatric tumors, with pediatric patients diagnosed with medulloblastoma treated with mostly, if not only, chemotherapy [2-4]. Sadly, barriers to the systemic route of administration limit drug penetration in the tumor microenvironment and include but are not limited to plasma protein binding, the blood-brain barrier, and structural and mechanical components within the tumor microenvironment all hindering the potential of therapeutic effect on the cancer cells [5-8]. In treating brain tumors, nearly 98% of small molecules and 100% of large molecules do not cross the blood-brain barrier, defining the blood-brain barrier as the most significant physical obstacle to treating brain tumors [9, 10]. For many chemotherapeutics such as doxorubicin (DOX), there are *in vitro* studies that show that the drugs are effective at inducing apoptosis of brain tumor cells, but due to the barriers to traditional administration, there are no consistent results *in vivo* [8, 11, 12]. These barriers require the use of high dosages to sustain therapeutic benefit at the tumor site, increasing the risk of long-term and short-term toxic side effects, especially to bystander organs including the heart and bone marrow [13-16]. Following current standard treatment, due to the long-term systemic effects of chemotherapy, medulloblastoma survivors are at considerable risk of neurological sequelae in cognition, fertility, heart and lung function, difficulty in school and work, lower physical fitness and function, as well as endocrine function [17]. Unfortunately, there is limited literature on designing drug delivery systems with hopes to overcome these challenges, presenting a significant gap in pediatric research.

Hydrogels are three-dimensional polymeric networks with the capability of sustaining high amounts of water and other biological fluids as well as being structurally engineered to control drug release kinetics [18-21]. Hydrogels can be designed to deliver drugs such as hydrophilic, small-molecule drugs, and macromolecular drugs beyond the blood-brain barrier by local administration as well as fine-tuning the size, architecture, and functionality of the polymer network of the hydrogel [22]. As drug delivery systems, they can achieve sustained and or controlled release drug delivery and provide beneficial outcomes by enhancing drug penetration while reducing drug toxicity when compared to conventional formulations [22, 23]. These drug-loaded hydrogels have been extensively investigated pre-clinically as a local drug delivery system for various applications [22, 24-26]. However, there is a lack of studies investigating hydrogels as a treatment method in pediatric oncology with a recent PubMed literature search of “*medulloblastoma hydrogels”* finding few primary studies [27-30].

To address the need for innovative drug delivery systems in pediatric oncology, we present a novel hydrogel drug delivery strategy designed to enhance drug penetration in treating medulloblastoma, the most common malignant pediatric brain tumor. A starch-based hydrogel designed primarily of amylopectin, with dimethyl sulfoxide (DMSO) as the solvent, was loaded with DOX. The starch-based hydrogel loaded with DOX described here can be tailored to hold large concentrations of drug, control drug release, and induce apoptosis of medulloblastoma cells. These results support the potential of local administration of hydrogel drug delivery systems as a method to overcome the challenging issues in pediatric oncology and contribute to advancements in developing strategies to enhance drug penetration while preventing exposure to non-target tissues.

## Materials and Methods

### Hydrogel Synthesis

Hydrogels were synthesized with a microwave-assisted organic synthesis method using an Anton Paar Microwave Reactor Monowave 200 (Graz, Austria). Microwave synthesis was chosen for its shortened reaction times compared to other methods [31], and preliminary tests, not presented here, were conducted to determine the optimal settings to synthesize hydrogel with the microwave reactor with a power during hold time of 150W, a hold time of 10 seconds, and a stir rate of 700 RPM. Three different concentrations of free-based DOX (MedKoo) were incorporated into hydrogels made with concentrations of no drug, 10 mg/mL DOX, and 100 mg/mL DOX. The hydrogel was synthesized around the drug in solution achieving 100% drug loading during the synthesis reaction. Freebase doxorubicin was added to a mixture of DMSO and amylopectin in the previously stated quantities and the sample was heating using the microwave reactor.

### HPLC Analysis

To assess drug release from the hydrogel system, a high-pressure liquid chromatography (HPLC) system, Agilent 1260 Infinity II Primary (Agilent Technologies USA), with a diode array detector equipped with a Poroshell 120 EC-C18 column of 2.7 μm particle size was used. The method developed (**Table 1**) consisted of a gradient elution with HPLC grade water with 0.1% trifluoretic acid and a solvent of 80% acetonitrile with 20% methanol. A 20 uL injection of each sample was eluted at a flow rate of 0.4 mL/min at 35°C with UV detection set at 254 nm. A wash vial containing the initial mobile phase was injected between samples. To quantify the concentration of samples being analyzed, a calibration curve was established with a stock concentration of DOX in DMSO and serially diluted with 1:2 ratio starting at 100 ug/uL.

**Table 1.** Method for HPLC Doxorubicin Analysis

| Method |  |
| --- | --- |
| Column | Poroshell 120 EC-C18, 50mm x 3mm, 2.7μm |
| Flow Rate | 0.4 mL/min |
| Column Temperature | 35C |
| UV | 254 nm |
| Injection Volume | 20μL |
| Needle Wash | [MeOH: ACN]:H2O = 1:1 |
| Solvents | A) Water with 0.1% TFA |
|  | B) 1:4 MeOH:Acetonitrile |
| Elution Conditions | Gradient |
|  | Time (min) |
|  | 0.0 |
|  | 17.00 |
|  | 18.00 |
|  | 20.00 |
|  | 24.00 |
|  | Stoptime: 13:00 min |
|  | Posttime: 2:00 min |
| | Poroshell 120 EC-C18, 50mm x 3mm, 2.7μm |

### Characterize Mechanical Rheology and Syringability Properties of Hydrogel

A rheometer was used to analyze the structure of the hydrogels synthesized at 3 wt%, 5 wt%, 10 wt%, and 20 wt% of amylopectin with either no drug, low-drug (10mg DOX), or high-drug (100mg DOX). To evaluate the architecture of the hydrogel, frequency sweep tests were performed to quantify the storage modulus, G’. In rheology, the storage modulus G’ (Pa) is used to describe the elastic properties represented by solid-state behavior. For the tests here, the storage modulus represents the energy needed for the material to flow into new conformations in response to applied stress. The storage modulus is proportional to the number of crosslinks or aggregates present in the polymer network.

The rheological measurements were performed at 25°C with an AR-2000 (TA Instrument, USA) with a 40 mm cone diameter and a cone angle of 1°. Samples were loaded with the same sample quantity avoiding overfilling and inadequate filling according to DIN 51810-1. Controlled sinusoidal strain test, where samples were sheared between two plates, were performed with the upper plate was moving back and forth over a stationary lower plate. To determine the appropriate amplitudes for all frequency sweep tests, the material’s linear viscoelastic region (LVR) was determined through controlled strain or controlled sweep tests. The LVR of an individual sample correlates to the region where tests can be performed independent of being imposed by stress and strain levels. This region was defined as the plateau value of G’. Oscillatory shear strain sweep tests were performed with a set angular frequency of 10 radians/second in a shear strain range of 0.01% to 100%. Oscillatory frequency sweep tests were performed at a fixed shear strain value of 1% in a range of 0.01 to 100 radians/second.

To test the injectable properties of the hydrogel, syringability tests were conducted with an Instron 5966 (Instron, USA) that applied an even force with a constant velocity to the plunger to extrude the hydrogel at a constant flow rate. While pressing, the Instron quantified the injection force required overtime to maintain a constant flow rate. The syringe was set in a stage, and parameters were set to test flow rates from syringe of 2 to 8 mL/h with a 1 mL syringe and 25G needle.

### Drug Release Profiles

Dissolution studies were performed by injecting hydrogel into the cell culture medium under appropriate sink conditions. Samples were collected at predetermined time intervals (0.5, 1, 2, 4, 6, 24, and 48 hours) and replaced. All drug release samples were stored in a cell culture incubator set at 37°C in a 5% CO_2_ humified atmosphere. HPLC was used to determine if drug structure remained intact upon release from the delivery system and to determine the correlation between the mesh size of the polymeric network and drug release kinetics. The schematic of the process used to analyze drug release is illustrated in **Fig. 1**. the amount of drug released from the hydrogel at a given time was quantified by HPLC [32].

**Figure 1.**
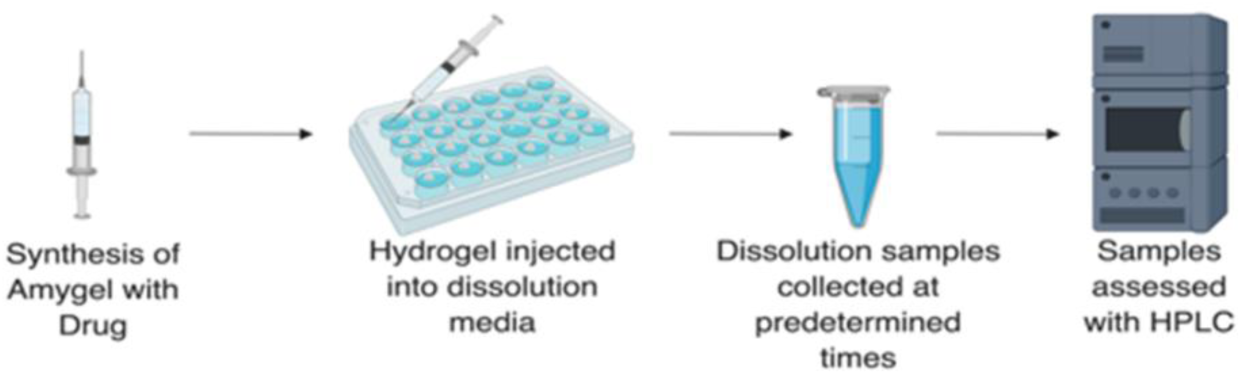
Methods schematic for quantifying durg realease from hydrogels over time. Created with Biorender.com

### Cell Culture

DAOY medulloblastoma cells (American Type Culture Collection, USA) were used for the *in vitro* studies. The DAOY cell line is the most cited and oldest medulloblastoma cell line and was selected for this study because of its strong transcriptional profiling data supporting its classification as an SHH cell line [32]. The cells were cultured in EMEM (Eagle’s Minimum Essential Medium; Sigma Aldrich, USA) supplemented with 10% FBS (fetal bovine serum; Sigma Aldrich, USA and 1% penicillin/streptomycin (Sigma Aldrich, USA) in T-75 culture flasks (Corning Inc., USA). The cells were cultured at 37°C in a 5% CO_2_ humified atmosphere. Media was replaced every 72 hours. For subculturing, trypsin (Sigma Aldrich, USA) was used to rinse and detach the cells. The cells in trypsin were incubated for 3-5 minutes at 37°C and visually inspected for detachment from the cell surface before adding the medium cocktail to deactivate the trypsin. The cells were split at 1:3 or 1:6 depending on confluency.

### *In Vitro* Cell Viability

A CellTiter 96® AQueous One Solution Cell Proliferation Assay (MTS assay; Promega Corporation, USA) was used to determine the number of viable cells in proliferation. This commonly used colorimetric method contains a tetrazolium compound [3 (4,5-dimethylthiazol-2-yl)-5-(3-carboxymethoxyphenyl)-2-(4-sulfophenyl)-2H-tetrazolium, inner salt; MTS] and PES (phenazine ethosulfate), an electron coupling reagent. To perform analysis, 3,000 cells per well were plated in 96-well plates for 24 hours before treatment to allow for adherence. Cells were then exposed to dissolution samples collected at different time intervals up to 24 hours. HPLC was used to determine the DOX concentration of the dissolution samples that were then added to the cells. The negative control consisted of cells not treated and the positive control consisted of cells treated with a stock solution of DOX. This method was modified from previous medulloblastoma studies and biomaterial response evaluations [33, 34]. The CellTiter 96® AQueous One Solution Reagent was added to the cell plates and incubated for 1-4 hours as indicated in the protocol. Absorbance at 490nm was measured using a microplate reader. Results from this study were used to determine the cytotoxicity response of the DAOY cell line to DOX released from the hydrogel.

### Statistical Analysis

Prism 9 (GraphPad, USA) was used to analyze the results. All experiments were performed in triplicate. All results depict mean values with error bars representing standard deviation. One-way ANOVA tests were performed to compare cells not treated (control), exposed to empty hydrogel, and drug-loaded hydrogel at each time point. For *in vitro* studies, a t-test was used to determine if there was a significant difference between cells exposed to empty hydrogels versus doxorubicin-loaded hydrogels.

## Results

### Hydrogel Synthesis Processes Leaves Doxorubicin Intact

For these experiments, three different amylopectin concentrations of hydrogels were synthesized: 3, 5, and 10 wt%. An HPLC method was developed, and a linear calibration curve was established to quantify the amount of doxorubicin and to assess drug structure and release behavior (**Fig. 2A**) with the equation for the area (mAU*sec) = 125.29792*ug/uL - 16.89752 with an R^2^ value of 0.99998. The peak of interest for detecting doxorubicin in solution was identified at 254 nm, consistent with prior studies [35-37]. For 0.1-100 ug/mL DOX in DMSO, a clean and distinctive peak was detected at the same time around 15 minutes, with the intensity of absorbance increasing with concentration (**Fig. 2**).

**Figure 2.**
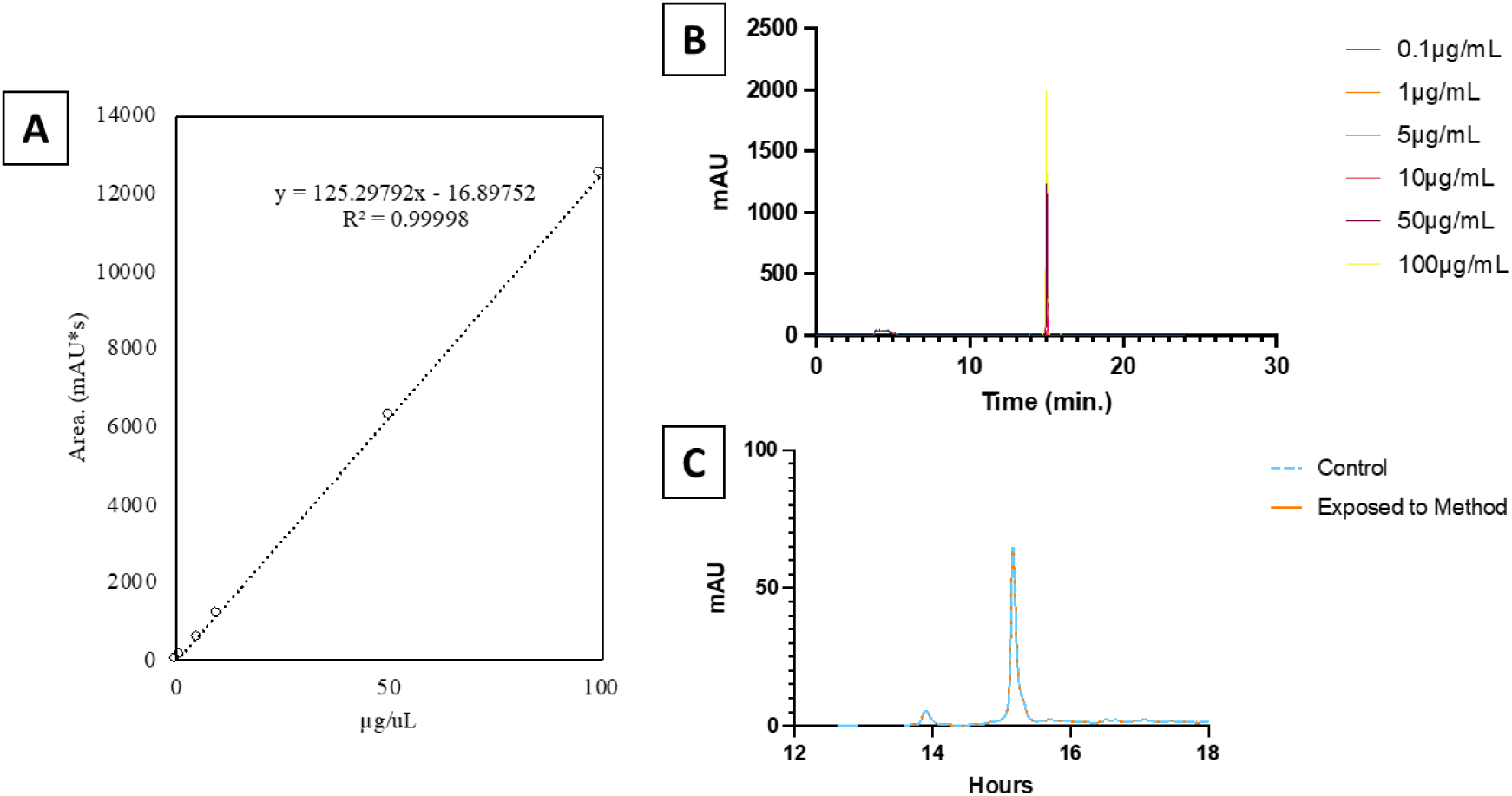
Doxorubicin Quality Verification After Gel Preparation. (A) Calibration curve of DOX used to quantify drug quantity in drug realese studies. (B) DOX HPLC reading was determined in unalteed drug for 0.1, 1, 5, 10, 50 and 100ug/mL concentrations and (C)

With this synthesis method, 100% drug loading was achieved with even distribution throughout the volume of gel. During method development, DOX was dissolved in DMSO and heated in the reactor then analyzed with HPLC to show that the drug was unaltered by the synthesis method. Chromatographic results confirmed a negligible shift in the peak identified as DOX between control and DOX released from the hydrogel (**Fig. 3**), confirming the synthesis method did not affect the drug’s structure.

**Figure 3.**
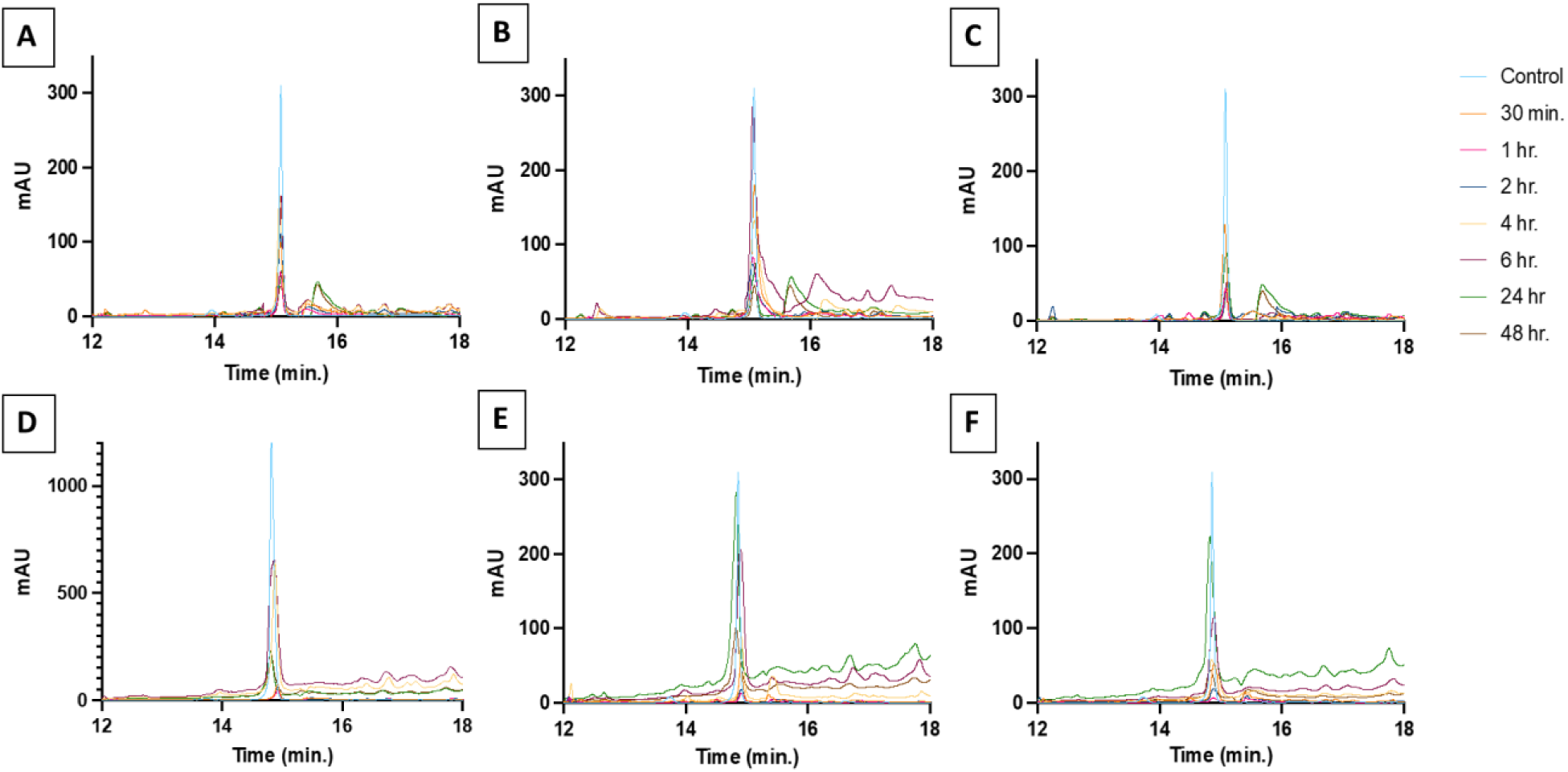
Chromatography of DOX Dissolution from Hydrogel. (A-C) low-dose DOX (10 mg) and (D-F) high-dose DOX after incubation for dissolution at various time points. Test conducted at 3 different concentrations of hydrogel: (A,D) 3 wt%, (B,E) 5 wt%, and (C,F) 10 wt%

### Drug Released at Higher Percentages from Lower Concentration Gels

The cumulative drug release (% loaded dose) of each group of hydrogels was quantified, and an initial burst release was evident in all hydrogel groups followed by sustained drug release over time (**Fig. 4**). At 6 hrs., drug release for hydrogels loaded with 10mg/mL of doxorubicin was 55.1%, 31.3%, and 15.9% respectively for 3, 5, and 10 wt% (**Fig. 4A**). At 6 hrs., drug release for hydrogels loaded with 100mg/mL of doxorubicin was 38.4%, 30.9%, and 21.4%% for respectively for 3, 5 and 10 wt% (**Fig. 4B**). At 48 hrs, the cumulative drug release (%) for hydrogels loaded with 10mg/mL doxorubicin was 61.3%, 45.5%, and 22.7% respectively for 3, 5, and 10 wt% (**Fig. 4C**). The average cumulative drug release (%) for triplicate hydrogels loaded with 100 mg/mL doxorubicin at 48 hrs. was 65.4%, 52.8%, and 36.8% respectively for 3, 5, and 10 wt% (**Fig. 4D**).

**Figure 4.**
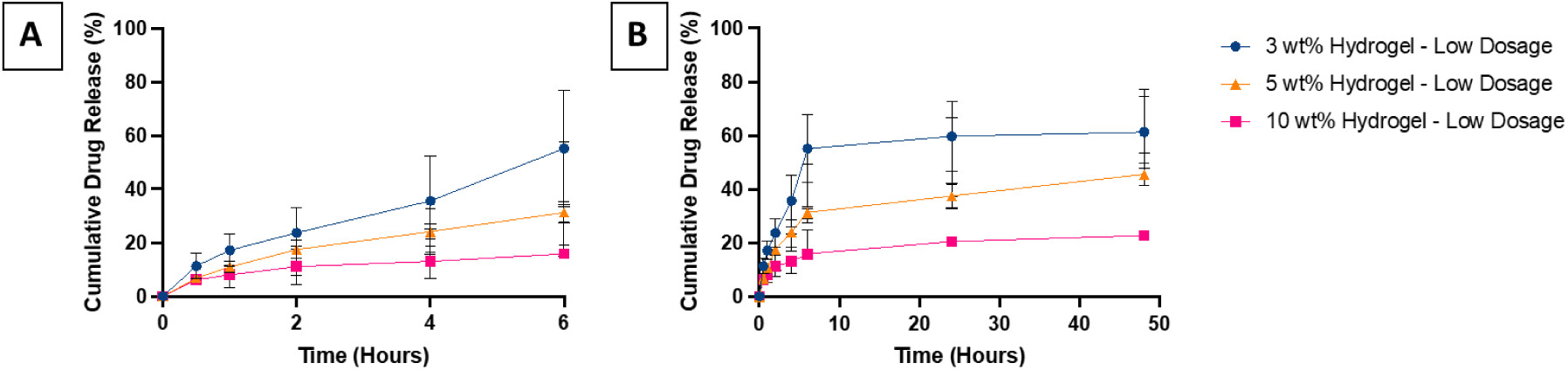
DOX Realsed from Hydrogel. DOX realese profile for 3, 5, and 10 wt% hydrogels over (A) the initial burst up to 6 hours and (B) until realesed plateaus at 48 hours low-dose construts.

### Higher Concentration Gels Have Increased Strength

Rheological tests were performed to compare the structure of the hydrogel compositions. Oscillatory shear sweep tests validated that 1% shear strain values are within the linear viscoelastic region of the hydrogel. Per ISO 6721-10 and EN/DIN EN 14770, the tolerance range of the plateau value of G’ is within +5% deviation at a strain of 1%.

The 3 wt% hydrogels analyzed here saw a linear increase in the storage modulus from 1-100 radians/second (**Fig. 5A**). In the 5 wt% hydrogels after the linear phase, there was a drop in the storage modulus of the low-dose hydrogels and an increase for the empty and high-dose hydrogels when angle frequencies increased above 100 radians/second (**Fig. 5B**). While the 10 wt% hydrogels saw an increase in variation between the samples, at increased angle frequencies the 10 and 20 wt% hydrogels continued to see a linear increase in the storage modulus (**Fig. 5C-D**).

**Figure 5.**
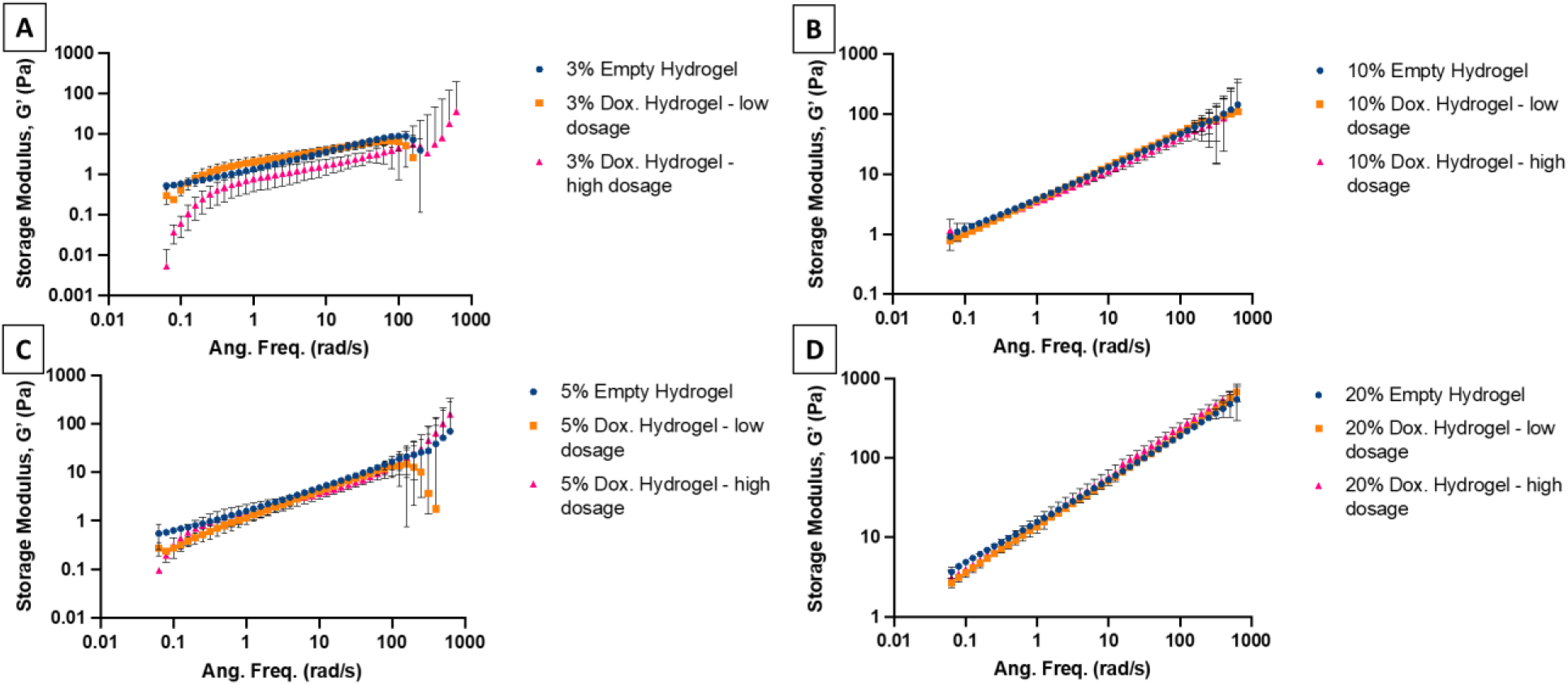
Rheology Determined Storage Modulus for DOX Loaded Hydrogels. Storage modulus (Pa) determined from the addition of an angular frequency of 0.1-1000 radians/sec on empty, low-dose, and high-dose DOX loaded hyrdrogels of (A) 3 wt%, (B) 5 wt%, (C) 10 wt%, and (D) 20 wt%.

3, 5, and 10 wt% hydrogels with and without DOX were tested for syringability by determining the injection force required to maintain three different flow rates (2, 5, and 8 mL/hr). For all the flow rates, the 3 wt% hydrogels required the least force, followed by 5 and 10 wt% (**Fig. 6**). Increasing flow rates from 2 to 8 mL/hr also resulted in increased injection force (**Fig. 6**). The empty 10 wt% hydrogels had a noticeably increased requirement compared to all the other hydrogels (**Fig. 6 C, F, I**).

**Figure 6.**
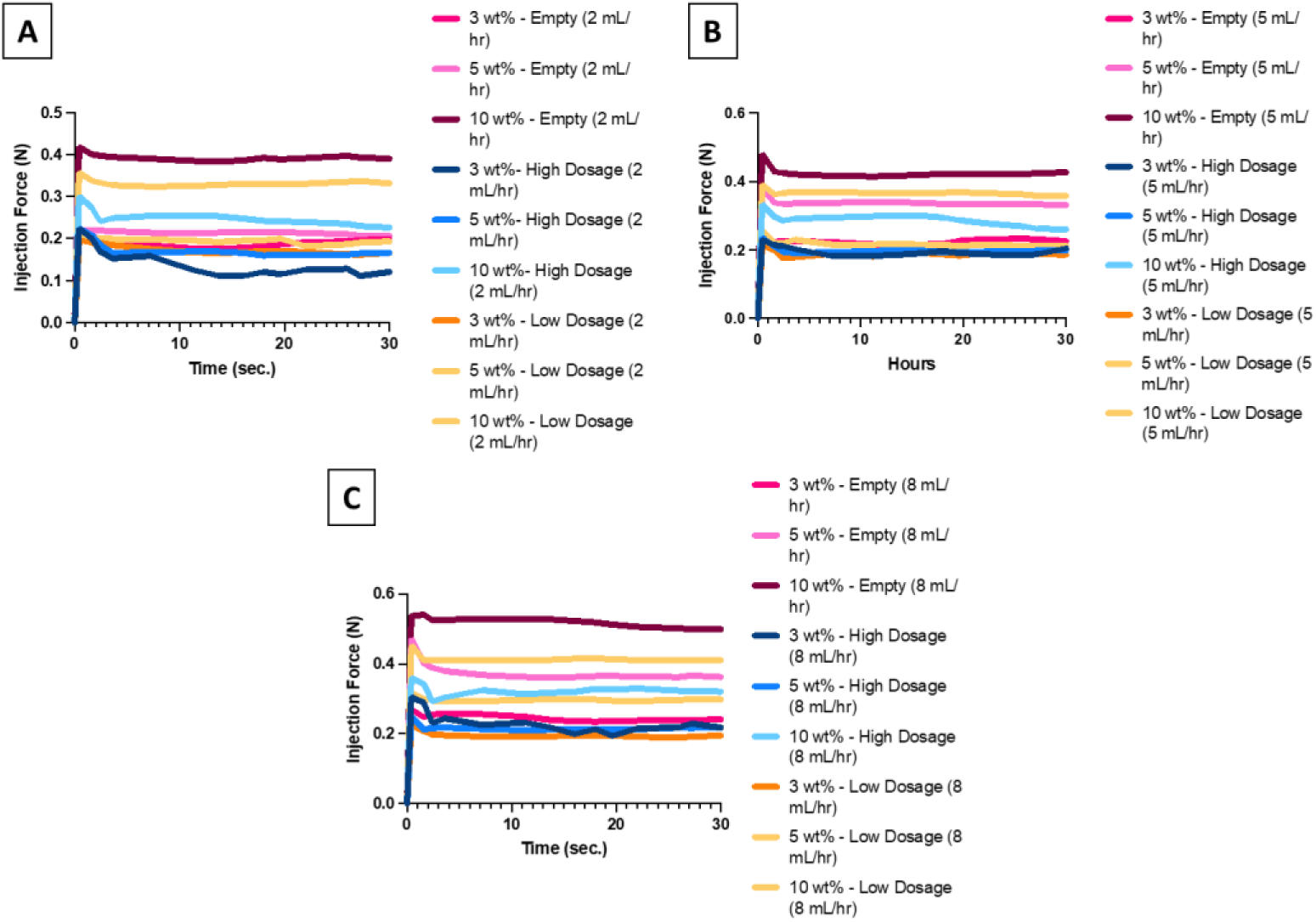
Syringability results of hydrogels across various concentration, dosages, and injection rates. Each graph includes gels made of 3 wt%, 5 wt%, and 10 wt% with no drug, high dose DOX, or low dose DOX. Figures represent the change in injection force needed to maintain an injection rate of (A) 2mL/hr, (B) 5 mL/hr, and (C) 8 mL/hr.

### DOX Released from Hydrogel Maintains *In Vitro* Cell Cytotoxicity

The DOX hydrogel drug delivery system was tested for *in vitro* cytotoxicity against the DAOY medulloblastoma cell line using an MTT assay. In general, DAOY medulloblastoma cells were found to be cytotoxically sensitive to both the DMSO solvent and the DOX (**Fig. 7**). There was a significant (p<0.05) drop in cell viability from the untreated controls and all the drug and empty hydrogel treatments (**Fig 7A**). There was an increase in cell viability in the empty hydrogels as the amylopectin concentration increased. Hydrogels loaded with doxorubicin showed significant cytotoxicity compared to control after 24 hours. These results support that DOX released from hydrogel is active and had a therapeutic effect. Regardless of the concentration of the hydrogel, there was no significant difference in cell viability across DOX-loaded hydrogels. In the 3 wt% hydrogels, there was no significant difference (p>0.05) in viability between the cells treated with DOX loaded and empty hydrogel (**Fig. 7B**). In the 5 and 10 wt% hydrogels, there was a significant difference (p<0.05) in viability between the cells treated DOX loaded and empty hydrogel (**Fig. 7C-D**).

**Figure 7.**
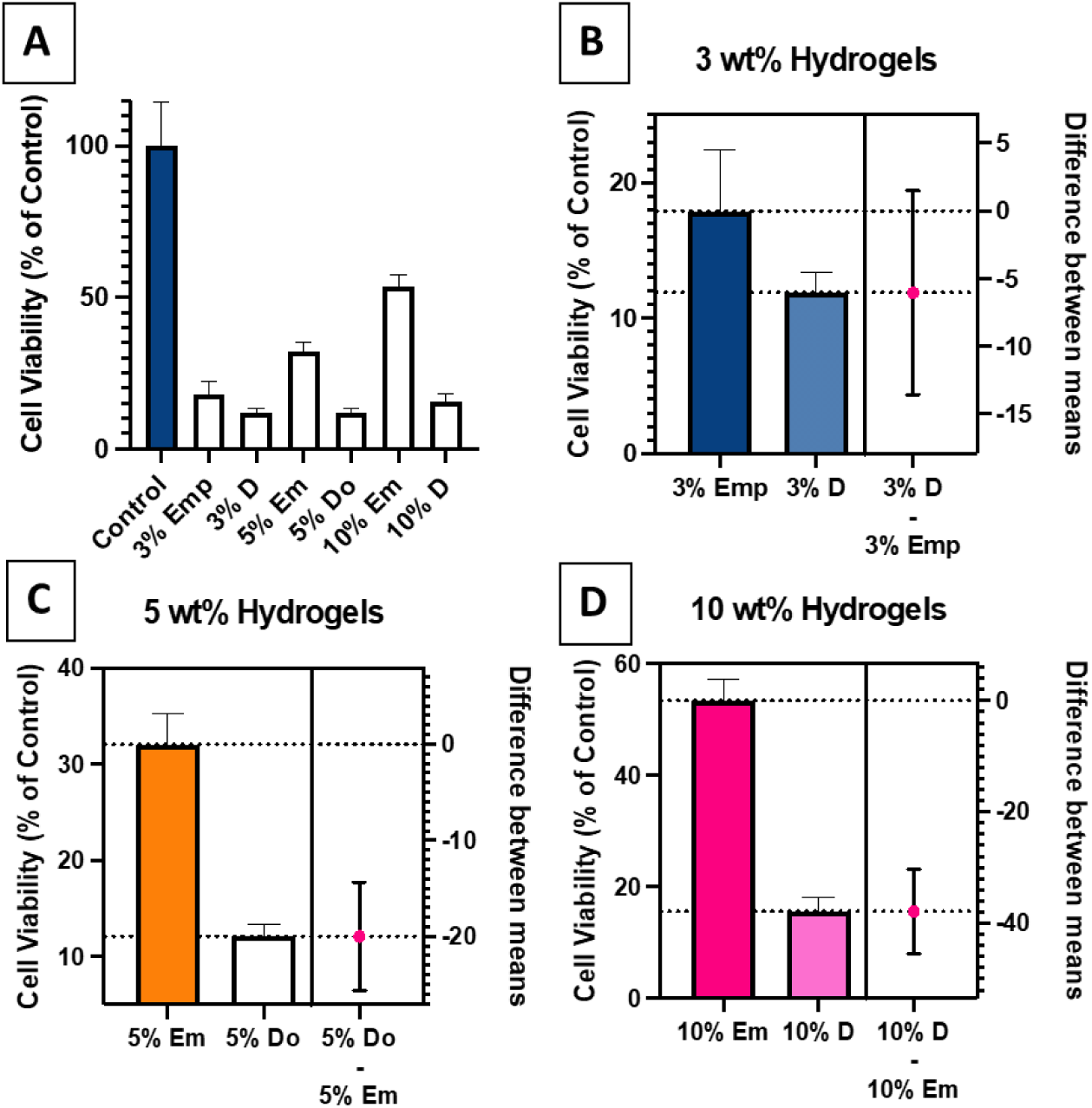
Hydrogel Released DOX is Cytotoxic *In Vitro*. (A) Cell viability results of DAOY cells after treatment with hydrogels depicting in-vitro response to hydrogels. Focused look at (B) 3 wt%, (C) 5 wt%, and (D) 10 wt% includes the difference between empty and DOX loaded hydrogel to isolate the effects of released DOX. * p<0.01

## Discussion

In this study, we showed that DOX can be incorporated into a novel amylopectin hydrogel network, maintain its structure, be released at moderate rates, and maintain cytotoxic efficacy against a medulloblastoma cell line. A significant challenge in designing drug delivery systems is the inability to load drugs into the system. Chemotherapy agents are often not soluble in water and are chemically altered to create aqueous intravenous formulations. DOX, the drug tested in this study, is known to be one of the most potent chemotherapy agents and is used to treat a multitude of tumors in adult and pediatric patients [38]. The primary mechanism of action of doxorubicin is intercalation within DNA base pairs, inducing breakage of DNA strands causing inhibition of RNA and DNA synthesis [38, 39]. Doxorubicin is associated with severe adverse reactions including cardiac toxicity, which is why there is a great need to reduce its systemic distribution. Therefore, by being able to construct a hydrogel that can deliver DOX locally to avoid systemic distribution and maintain a steady release of active drug, the current hydrogel could minimize the negative effects of DOX therapy and maintain its local efficacy.

For the current hydrogel, we used amylopectin, a highly branched carbohydrate component of starch, because of its ability to be thermally gelatinized to form a network and its branched tree-like topology that allows for more bonding compared to other polysaccharides used in the development of hydrogels like chitosan, dextran, and hydroxyethyl cellulose [40, 41]. An additional advantage of using a starch-based hydrogel compared to other local drug delivery systems is that the matrix can be designed with drug encapsulated and be safely and readily injected as seen in previous *in vivo* studies [40]. The three-dimensional network made of crosslinked polymer chains allows for drugs to be integrated into and diffuse out in a manner that can be controlled by adjusting the mesh size [22]. DOX’s cytotoxic mechanism of action involves forming complexes with DNA by intercalating between base pairs, inhibiting cell replication [38, 39]. Therefore, it was theorized that DOX could similarly disrupt the polymer network, weakening the structure of the hydrogel. However, this pattern was only subtly seen in 5% and 10% hydrogels in this study even when doxorubicin concentration reached 100 mg/mL (**Fig. 3**). The formation of aggregates was still present in creating the hydrogel network as represented in the rheology data which further showed that an increase in amylopectin concentration allowed for increased network stability (**Fig. 5**). For releasing the DOX, the rate was observed to be inversely proportional to polymer concentration (**Fig. 4**). The hydrogels made with 3 wt% had the highest burst release and fastest drug release over 48 hours, while the 10 wt% hydrogels showed the least burst release and drug released over the 48 hours. These results confirm drug release can be controlled by hydrogel composition, with the injectable hydrogel designed for this study allowing for a high dose of 1 mg of doxorubicin delivered in a single injection volume of 10 uL.

From the rheology results (**Fig. 5**), it was shown that higher polymer concentration is associated with increased storage modulus, contributing to the increased injection force (**Fig. 6**). Mechanical testing results also quantified the initial break force and sustaining (gliding) force for each of the hydrogels. According to the SS-EN ISO 11608-3, which is an international standard for needle-based injection systems for medical use including medical devices used for pharmaceutical administration, the initial break force should not exceed 40 N nor the gliding force exceed 20 N. For all the samples tested, this drug delivery system meets those requirements. The rheological results further support the ability to control network mesh size in designing a hydrogel as a drug delivery system. Depending on the drug release profile needed from the network, amylopectin concentration can be adjusted to control the dose administered to the tissue.

DMSO was used as the solvent to form the hydrogel in this study. Two of the advantageous properties of DMSO are enhanced tissue penetration and high drug solubility [42-44]. Here, DMSO was used to maximize local drug concentration and to enhance penetration in the tumor microenvironment. However, medulloblastoma and specifically the DAOY cell line is sensitive to DMSO. The results demonstrate that an injectable polysaccharide hydrogel formulated with the tissue penetrating agent, DMSO, has high DOX loading properties, can achieve 100% drug loading when synthesized using the method described, releases DOX in its active form, and achieves sustained release over time. A study is currently underway to test this formulation *in vivo* using a nude mouse model and the same DAOY cell line.

